# High-order brain interactions distinguish wakefulness, anaesthesia, and recovery induced by deep brain stimulation

**DOI:** 10.64898/2026.06.01.728390

**Authors:** Camilo Espinosa-Curilem, Lynn Uhrig, Jordie Tasserie, Ruben Herzog, Marilyn Gatica, Andrea Luppi, Jorge F. Silva, Bechir Jarraya, Rodrigo Cofre

## Abstract

Understanding how consciousness depends on large-scale brain interactions is key for both the neuroscience of consciousness and clinical translation. However, it requires moving beyond classical pairwise descriptions of functional connectivity, which cannot capture the collective dependencies emerging across multiple brain regions. Here, we use multivariate information theory measures to characterize how higher-order interactions re-organize across states of consciousness. Specifically, we apply O-information to resting-state fMRI data from non-human primates to quantify whether multiregional brain dynamics are dominated by synergistic or redundant information sharing. We analyse two complementary datasets: (i) wakefulness and anaesthesia-induced loss of consciousness using different molecular agents (propofol, sevoflurane, ketamine), and (ii) the recovery of consciousness driven by central thalamic deep brain stimulation during propofol anaesthesia, indexed by behavioural responsiveness. We identify optimal regional subsets whose O-information robustly discriminates conscious from non-responsive states under two complementary optimization polarities. The first captures elevated redundancy in conscious scans that decreases under anaesthesia, providing robust discrimination and placing high-voltage central-thalamus stimulation closer to wakefulness. The second captures a synergy-to-redundancy transition, prominent in multi-anaesthesia conditions but context-dependent across datasets. Discrimination performance depends on interaction order: redundancy-based signatures improve with increasing subset size, whilst synergy-based signatures peak at low orders. Higher-order informational features significantly outperform pairwise functional connectivity, particularly for synergistic signatures which remain invisible to correlations. These findings demonstrate that consciousness is reflected in the reconfiguration of higher-order interaction structures, with distinct informational substrates requiring multivariate characterization beyond pair-wise connectivity.

## I. Introduction

Understanding how conscious awareness emerges from large-scale brain activity remains a central challenge in systems neuroscience. A key difficulty lies in identifying neural signatures of consciousness that are robust across distinct physiological and pharmacological manipulations, yet specific to the presence or absence of behavioural responsiveness. Historically, attempts to address this problem have been shaped by localisationist views. While these ideas were gradually replaced by connectionist and, more recently, network-based frameworks [1], [2], much of contemporary neuroscience still relies on descriptions of brain organization grounded in pairwise interactions between regions [3], [4].

Network neuroscience has undoubtedly transformed our understanding of brain function, revealing how cognition and perception arise from coordinated activity across distributed systems [3], [4]. However, pairwise functional connectivity (FC) provides only a partial description of neural dynamics. By construction, it cannot capture higher-order interactions (HOIs), collective dependencies among triads or larger sets of regions, that are increasingly recognized as fundamental to multivariate neural computation. Growing evidence from neuroimaging and computational modeling indicates that such HOIs play a critical role in shaping brain dynamics, motivating the rapid development of analytical frameworks based on hypergraphs [5], [6], simplicial complexes [7], [8], and multivariate information theory [9], [10], [11], [12]. These approaches offer principled tools to characterize how information is jointly processed by groups of regions, beyond what can be inferred from pairwise correlations alone.

HOIs are particularly relevant in the study of consciousness. Conscious brain activity is thought to rely on a delicate balance between information integration (binding distributed signals into unified percepts) and information segregation (preserving specialized, local processing) [13], [14], [15]. This integration–segregation framework, originally articulated by Tononi, Sporns, and Edelman [16], posits that consciousness emerges neither from complete independence nor from global synchrony, but from an intermediate regime of structured coordination across multiple spatial scales. Disruptions of this balance, including general anaesthesia, disorders of consciousness, and targeted neuromodulation, reliably alter behavioural responsiveness [17], [18], [19], [20]. Yet, identifying multivariable neural features that consistently track these state transitions remains an open challenge.

Studies in humans and non-human primates (NHPs) have shown that loss of consciousness under general anaesthesia is associated with reduced global integration, increased modularity, and constrained, anatomically driven FC patterns [21], [22], [23], [24], [12]. While these findings have provided important insights, they are based on pairwise FC measures [3], [25], [26], which are inherently limited in their ability to capture collective, multivariate dependencies [11], [27], [10], [28]. Crucially, anaesthetic agents such as propofol, sevoflurane, and ketamine act through distinct molecular and circuit-level mechanisms, yet converge phenomenologically on a common endpoint: a state of generalized unresponsiveness. This raises a fundamental question: are there sets of brain regions whose higher-order informational interactions reliably differentiate awake from unresponsive states? And if so, are these signatures conserved independently of the specific anaesthetic mechanism?

Information-theoretic measures provide a natural framework to address this question. Multivariate decompositions reveal that neural systems can process information synergistically (where information is present only in joint activity, but not in the parts) or redundantly (where information is duplicated across regions) [9], [29]. Synergistic and redundant interactions have been demonstrated to differentiate distinct brain states in fMRI studies [10], [30], [31], [32], [11], [27].

A particularly powerful measure in this context is O-information (Ω) [9], which quantifies the balance between redundancy (Ω > 0) and synergy (Ω < 0) within a set of three or more variables. It captures the degree to which the collective behavior of a group of variables is better described by shared, overlapping information (redundancy) or by dependencies that only emerge when all variables are considered together (synergy). Unlike pairwise metrics, O-information explicitly captures collective interdependencies and enables the identification of sets of regions whose joint dynamics are informative of global brain states [11]. Although existing studies have characterized important insights into information dynamics under anaesthesia [33], [34], [12], [31], whether O-information signatures of consciousness are conserved across distinct anesthetic agents and whether they can be restored through neuromodulation remain open questions.

Recent methodological advances, including computational toolboxes such as THOI [35], JIDT [36], and the HOI toolbox [37], have made it increasingly feasible to apply multivariate information-theoretic measures to large-scale fMRI datasets.

In this work, we apply multivariate information theory to characterize HOIs associated with conscious and unresponsive states in NHPs. We analyse two complementary fMRI datasets (Fig. 1A). First, a multi-anaesthesia (MA) dataset in which non-human primates were scanned during wakefulness and under three distinct anaesthetic agents (propofol, sevoflurane, and ketamine). Second, a deep brain stimulation (DBS) dataset in which non-human primates were scanned during wakefulness, under propofol anaesthesia without stimulation, during stimulation of a control site (ventrolateral thalamus, VT), and during recovery of behavioural responsiveness induced by central thalamic stimulation at two intensities [20].

**Fig. 1.**
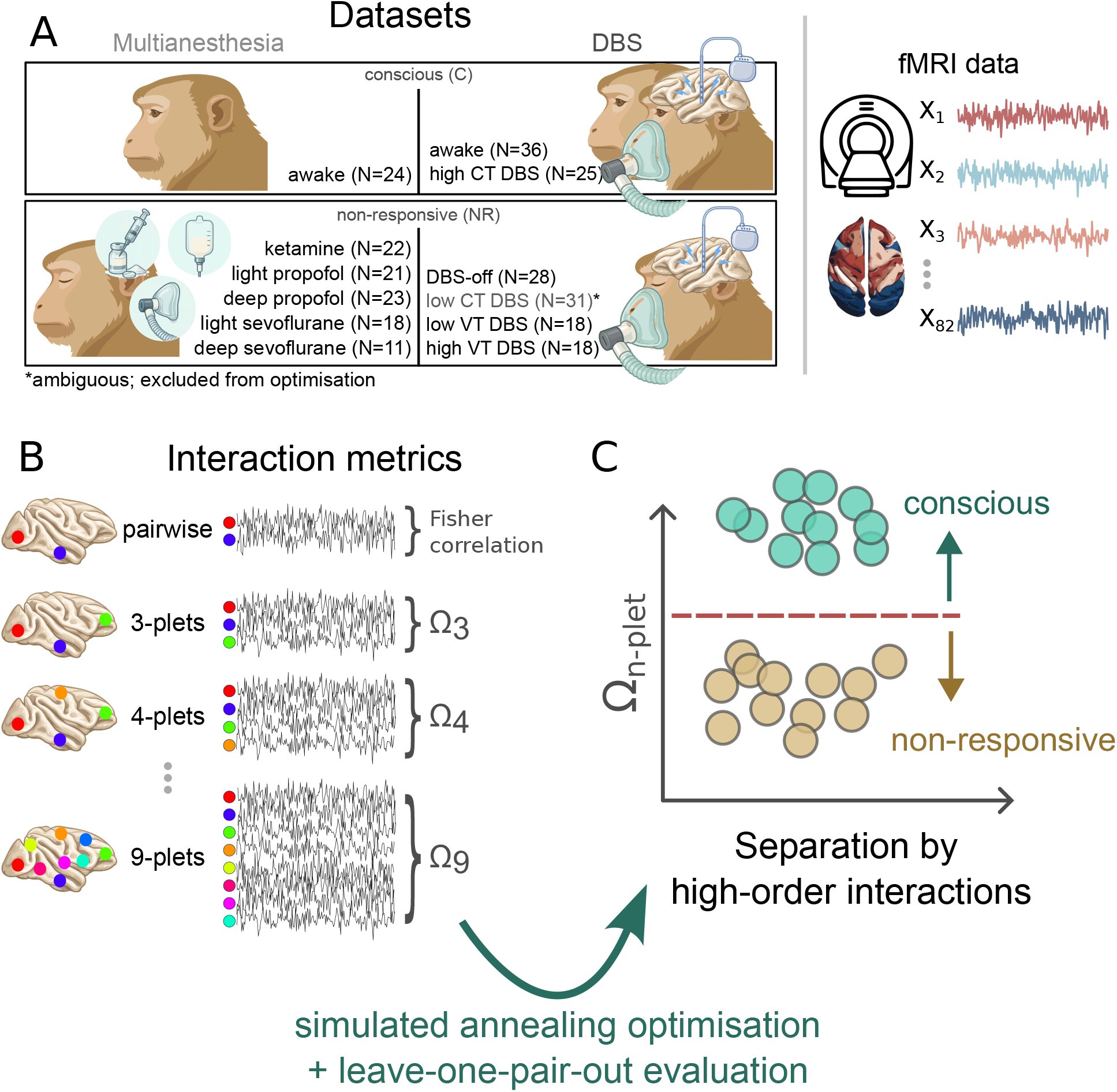
Estimating and comparing higher-order interactions across states of consciousness. **A:** Two complementary datasets. The multianaesthesia (MA) dataset includes awake scans and multiple anesthetic agents (ketamine, propofol, sevoflurane at varying depths). The deep brain stimulation (DBS) dataset includes awake scans, anesthetized scans, and central thalamic stimulation at different intensities (low-intensity stimulation marked as ambiguous and excluded from optimization). For both datasets, whole-brain resting-state fMRI (rs-fMRI) is acquired across all conditions. **B:** Computation of interaction metrics. Regional BOLD time series are extracted and analyzed in subsets of *n* regions (“*n*-plets”). For each *n*-plet size (*n* = 3 to *n* = 9), we compute O-information (Ω) that quantifies the balance between synergistic and redundant interactions across the *n* regions and a pairwise baseline metric. **C:** Identification of optimal discriminative subsets. Using simulated annealing with leave-one-pair-out cross-validation, we identify *n*-plets whose O-information (Ω_*n*-plet_) best separates conscious (C) from non-responsive (NR) states, with the aim of characterising how higher-order interactions change across conditions.

This dual-dataset design allows us to address three core questions: (i) whether higher-order informational signatures of unresponsiveness are conserved across anaesthetic agents with distinct mechanisms of action; (ii) whether specific sets of brain regions optimally discriminate awake from unresponsive states; and (iii) whether neuromodulatory interventions that restore behavioural responsiveness also restore higher-order informational signatures of wakefulness.

## II. Results

We first tested whether HOI signatures captured by Ω reliably differentiate conscious (C) from non-responsive (NR) scans using small subsets of brain regions (*n*-plets, Figure 1). Then, we checked whether this separation is conserved across distinct anaesthetic agents in the MA dataset, whether central-thalamus stimulation at high voltage in the DBS dataset is accompanied by Ω patterns closer to wakefulness, and characterised where in the brain the most discriminative *n*-plets are located. Next, we examined how these signatures vary with interaction order and compared higher-order features against a pairwise functional connectivity baseline (obtained by averaging all pairwise correlations within each *n*-plet). Finally, we assessed the generalisability of these signatures by testing whether optimal *n*-plets identified in one dataset could successfully discriminate conscious from non-responsive states in the other dataset.

### A. Higher-order signatures robustly discriminate conscious from non-responsive states

We defined two macrostates based on behavioural responsiveness. In the MA dataset, the conscious macrostate (C) corresponds to awake scans (24 runs), whereas the non-responsive macrostate (NR) includes all anaesthetised conditions (95 runs in total: ketamine, light and deep propofol, and light and deep sevoflurane). In the DBS dataset, C comprises non-anaesthetised wakefulness and high-voltage central-thalamus stimulation (61 runs), whilst NR includes anaesthetised scans without stimulation and ventral-thalamus control stimulation (64 runs). The intermediate condition of low central-thalamus stimulation (31 runs) was treated as ambiguous and excluded from optimisation, but is shown in Figure 2 for reference.

**Fig. 2.**
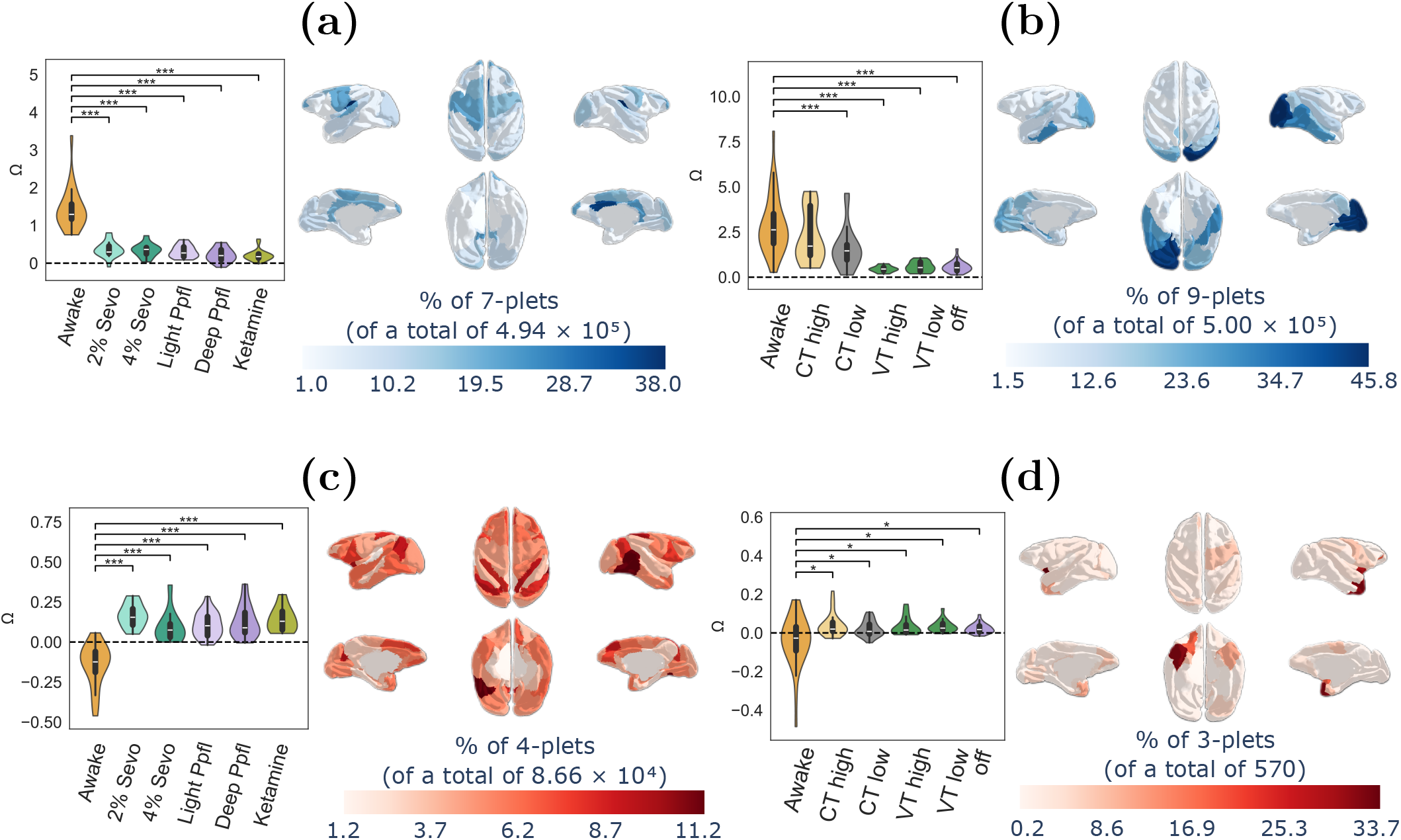
Non-responsiveness reconfigures the balance between redundancy and synergy in higher-order brain interactions. Violin plots show O-information (Ω) distributions across scans, evaluated on optimal discriminative *n*-plets for each dataset and optimisation polarity Ω_C_ > Ω_NR_ and Ω_NR_ > Ω_C_. Brain maps depict the relative frequency with which each region appears in high-performing *n*-plets of the same size. **(a, c)** Multi-anaesthesia dataset under Ω_C_ > Ω_NR_ (blue in brain maps) and Ω_NR_ > Ω_C_ (red in brain maps) optimisation. **(b, d)** Same as **(a, c)** for the DBS dataset. Low central-thalamus stimulation (excluded from optimisation) is shown for reference (CT low). Asterisks indicate Bonferroni-corrected Mann-Whitney test significance: ^∗^*p* < 0.05, ^∗∗^*p* < 0.01, ^∗∗∗^*p* < 0.001.

We then applied a simulated annealing procedure to identify, for each *n*-plet size *n* ∈ {3, …, 9}, the sets of regions whose Ω best separates C and NR scans, evaluated in a leave-one-pair-out (LOPO) scheme (see Methods). We considered two optimisation polarities that differ in the direction of the Ω contrast. In the Ω_C_ > Ω_NR_ context, we searched for *n*-plets with higher Ω in conscious scans by maximising ΔΩ = Ω_C_ − Ω_NR_.

Since Ω > 0 indicates redundancy-dominated interactions and Ω < 0 indicates synergy-dominated interactions, this polarity preferentially identifies interactions that are more redundancy-dominated (or less synergy-dominated) in C than in NR. Conversely, in the Ω_NR_ > Ω_C_ context we searched for *n*-plets with higher Ω in NR scans by maximising ΔΩ = Ω_NR_ − Ω_C_, thereby prioritising interactions shifting towards greater redundancy in the NR macrostate. Classification separability was quantified using the area under the precision-recall curve (PR-AUC) for classifying C versus NR scans, where PR-AUC = 1 indicates perfect separability (see Section V-D1).

Across the tested interaction orders *n* ∈ {3, …, 9}, we selected for each dataset and polarity the single *n*-plet with the highest PR-AUC. In the MA dataset, the best-performing Ω_C_ > Ω_NR_ *n*-plet included 7 regions and achieved PR-AUC = 0.996, whilst the best-performing Ω_NR_ > Ω_C_ *n*-plet comprised 4 regions and reached PR-AUC = 0.970. In the DBS dataset, the best-performing Ω_C_ > Ω_NR_ *n*-plet included 9 regions and attained PR-AUC = 0.974, whereas the best-performing Ω_NR_ > Ω_C_ *n*-plet comprised 3 regions with PR-AUC = 0.724. The regions included in each optimal *n*-plet are reported in Table II.

Figure 2 shows the distribution of Ω values across scans for each condition, evaluated on the corresponding optimal *n*-plet. Brain maps summarise regional involvement amongst high-performing *n*-plets at the same size as each optimum, showing which regions recurrently participate in discriminative higher-order interactions (see Section V-E for details). For each dataset and polarity, Ω distributions for NR scans were compared against the C condition using two-sided Mann-Whitney U tests with Bonferroni correction for multiple comparisons. Statistically significant differences are indicated by asterisks (exact values reported in Supplementary Tables III and IV).

In the MA dataset under Ω_C_ > Ω_NR_ optimisation (*n* = 7; Figure 2a), conscious scans exhibited substantially larger Ω (median Ω_C_ = 1.29) than all anaesthetic conditions (median Ω_NR_ = 0.25, *p* < 10^−3^), with practically no overlap between macrostates (PR-AUC = 0.996). Both conscious and non-responsive states showed predominantly positive Ω values (redundancy-dominated interactions), but wakefulness was characterised by significantly higher redundancy. Regional participation was concentrated in frontal and cingulate cortices, with the most frequent region appearing in 37.9% of high-performing *n*-plets. Under Ω_NR_ > Ω_C_ optimisation (*n* = 4; Figure 2c), awake scans clustered at negative Ω (median Ω_C_ = −0.12), whilst all anaesthetised conditions shifted to predominantly positive values (median Ω_NR_ = 0.13, *p* < 10^−3^), producing minimal overlap between macrostates (PR-AUC = 0.970). The corresponding regional participation was more widespread, spanning most of the cortex with comparatively stronger involvement in visual and motor cortices, whilst the most frequent region appeared in only 11.2% of high-performing *n*-plets. Notably, the separation between wakefulness and anaesthesia was conserved across all anaesthetic agents in both polarities.

In the DBS dataset under Ω_C_ > Ω_NR_ optimisation (*n* = 9; Figure 2b), Ω was elevated in conscious conditions (median Ω_C_ = 2.29), with awake and high-voltage central-thalamus stimulation showing larger values than anaesthetised and control-stimulation scans (median Ω_NR_ = 0.51, *p* < 10^−3^), achieving PR-AUC = 0.974. Both macrostates showed positive Ω values, with conscious states exhibiting higher redundancy. Regional participation was dominated by visual cortex, where the most frequent region appeared in 45.8% of high-performing *n*-plets. Low-voltage central-thalamus stimulation exhibited intermediate behaviour, with Ω values higher than NR scans yet still significantly different from wakefulness. Under Ω_NR_ > Ω_C_ optimisation (*n* = 3; Figure 2d), separability was substantially weaker (PR-AUC = 0.724). Awake scans spanned both signs of Ω with median near zero (median Ω_C_ = 0.0051), and both high- and low-voltage central-thalamus stimulation resembled NR conditions with mostly positive values (median Ω_NR_ = 0.0197, *p* = 1.79 × 10^−2^). Regional involvement was more localised than in the MA dataset, concentrated in right temporal cortex with peak frequency reaching 33.7% of high-performing *n*-plets. Notably, this context yielded substantially fewer filtered *n*-plets satisfying ΔΩ = Ω_NR_ − Ω_C_ > 0 (570 *n*-plets), consistent with the reduced robustness of this polarity in the DBS dataset.

### B. Higher-order interactions outperform pairwise connectivity with polarity-dependent order effects

Having identified optimal *n*-plets for macrostate discrimination, we next examined how separability depends on interaction order *n*. Using the optimal *n*-plets obtained from the optimisation procedure, we analysed each order *n* ∈ {3, …, 9} separately. For each dataset and polarity, we evaluated Ω for each optimal *n*-plet on every scan to obtain the order-dependent distributions shown in Figure 3a-d, and tracked separability across orders in Figure 3e,f using these higher-order features alongside a pairwise functional connectivity baseline.

**Fig. 3.**
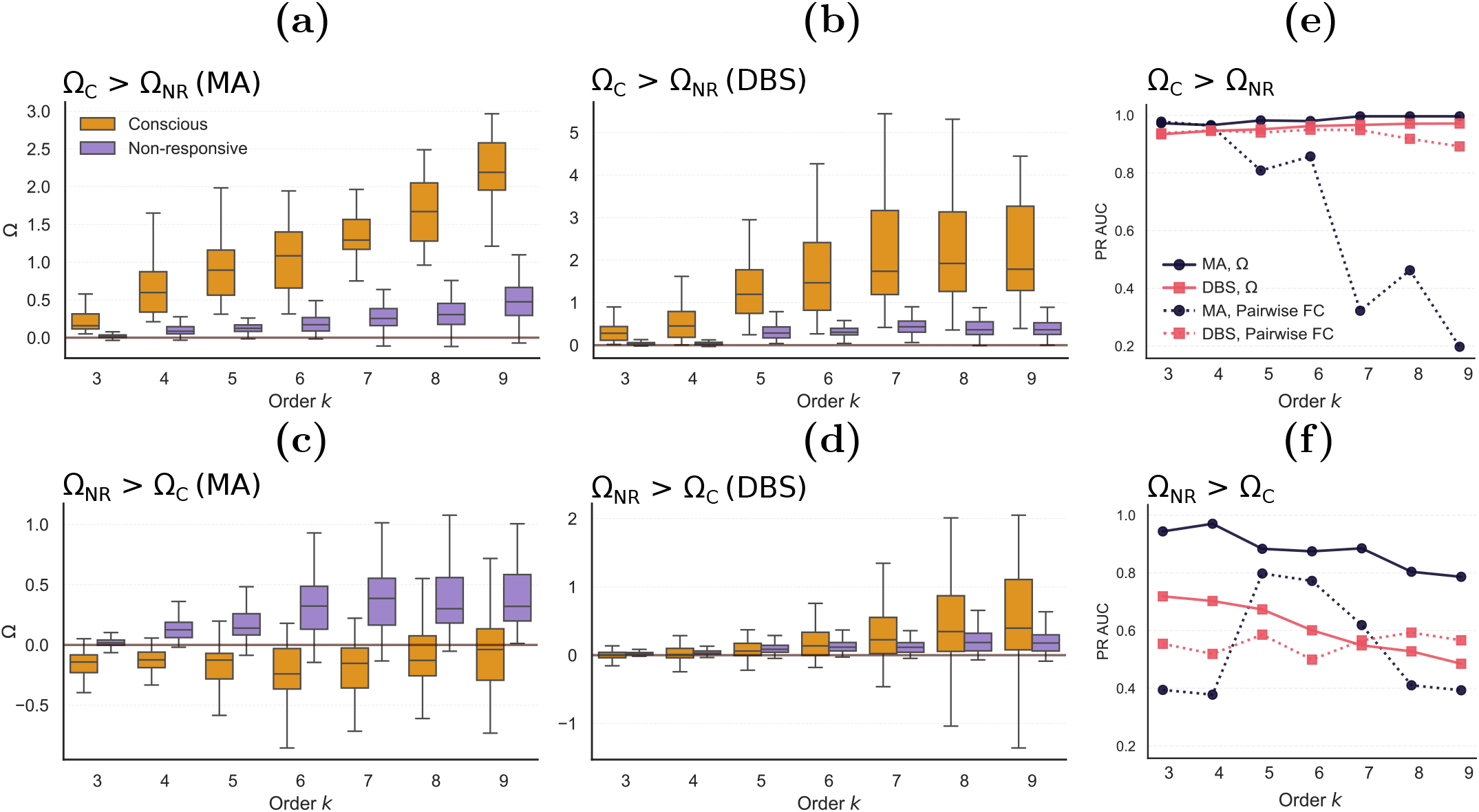
Macrostate separability shows polarity-dependent order effects and higher-order features outperform pairwise interactions. **(a**,**b)** Distributions of Ω for *n*-plets selected under Ω_C_ > Ω_NR_ optimisation in the multi-anaesthesia (MA) and deep brain stimulation (DBS) datasets, respectively. For each order *n* (3-9), we selected the best *n*-plet at that order according to PR-AUC and pooled Ω values across all scans. Boxplots summarise scan-wise values for conscious (C) and non-responsive (NR) scans. **(c**,**d)** Analogous distributions for Ω_NR_ > Ω_C_ optimisation. **(e**,**f)** PR-AUC for classifying C versus NR using HOIs (Ω, solid lines) and a pairwise functional connectivity baseline (mean Fisher *z*-values within the same *n*-plet, dashed lines), for Ω_C_ > Ω_NR_ (e) and Ω_NR_ > Ω_C_ (f) contexts.

For *n*-plets selected under Ω_C_ > Ω_NR_ optimisation (Figure 3a,b), Ω increased monotonically with *n* in both datasets and remained consistently larger in C than in NR. Although Ω in NR scans also increased with *n*, the separation between macrostates was generally larger at higher orders. This pattern translated into progressively improved discrimination performance (Figure 3e): PR-AUC increased monotonically with interaction order in both datasets, approaching near-perfect separability at the highest orders. The pairwise functional connectivity baseline (mean Fisher *z*-transformed correlation within each *n*-plet; see Section V-C) showed markedly different behaviour. In the MA dataset, pairwise features performed comparably to Ω at low orders but then declined dramatically at higher orders, whilst in the DBS dataset pairwise features remained relatively stable across orders but failed to improve with increasing order as Ω-based features did.

The Ω_NR_ > Ω_C_ optimisation behaved differently (Figure 3c,d). In the MA dataset, Ω was lower in C than in NR at low orders, as expected from the optimisation objective, with conscious scans showing predominantly negative values (synergy-dominated) and non-responsive scans showing positive values (redundancy-dominated). However, from *n* ≥ 5 onwards, Ω values became positive in both macrostates and the distributions increasingly overlapped, which appears to show that the synergy-to-redundancy transition dissolves at higher orders even among optimally selected regional subsets (which maximise ΔΩ at each order). In the DBS dataset, the contrast between C and NR was substantially weaker across all orders, with awake scans spanning both signs of Ω and showing considerable overlap with NR conditions even at low orders.

Correspondingly, discrimination performance (Figure 3f) showed opposite trends to the Ω_C_ > Ω_NR_ context. In the MA dataset, Ω-based PR-AUC peaked at low orders, achieving its maximum at *n* = 4 (PR-AUC = 0.970, the overall optimum reported in the previous section), then declined progressively with increasing order. In the DBS dataset, performance was lower overall and decreased monotonically with order from *n* = 3 (PR-AUC = 0.724, the overall optimum). Notably, pair-wise features performed substantially worse than Ω across all orders in both datasets. In the MA dataset, pairwise PR-AUC remained consistently low with no systematic trend across orders, whilst in the DBS dataset pairwise features showed no consistent pattern across orders. The marked superiority of Ω over pairwise correlations in this context demonstrates that the synergy-to-redundancy transition captured by Ω_NR_ > Ω_C_ *n*-plets is fundamentally invisible to pairwise connectivity measures.

### C. Cross-dataset generalisability of higher-order signatures

Having established the four optimal *n*-plets (one per dataset and polarity) and their within-dataset discrimination performance, we next asked whether these signatures generalise across experimental contexts. To this end, we directly applied each *n*-plet to scans from the other dataset and from both datasets pooled together, without any additional optimisation, using the same Ω values and classification scheme (Table I, Fig. 4).

**TABLE I.** Cross-dataset generalisability of higher-order signatures. Optimal *n*-plets identified in one dataset were tested on the other dataset and on both combined to assess transferability of higher-order informational signatures across experimental contexts.

| Origin Dataset | Evaluated Dataset | Polarity | $n$-plet size | PR-AUC |
| --- | --- | --- | --- | --- |
| MA | DBS | $\Omega_C > \Omega_{NR}$ | 7 | **0.915** |
| MA | DBS | $\Omega_{NR} > \Omega_C$ | 4 | **0.832** |
| DBS | MA | $\Omega_C > \Omega_{NR}$ | 9 | 0.212 |
| DBS | MA | $\Omega_{NR} > \Omega_C$ | 3 | 0.312 |
| MA | MA+DBS | $\Omega_C > \Omega_{NR}$ | 7 | **0.873** |
| MA | MA+DBS | $\Omega_{NR} > \Omega_C$ | 4 | 0.313 |
| DBS | MA+DBS | $\Omega_C > \Omega_{NR}$ | 9 | **0.801** |
| DBS | MA+DBS | $\Omega_{NR} > \Omega_C$ | 3 | 0.351 |

**TABLE II.** Optimal *n*-plets identified in the MA and DBS datasets and used in Fig. 2. Region indices correspond to the Regional Map parcellation described in Methods. PR-AUC values quantify C–NR separability for each *n*-plet.

| Dataset | Polarity | PR-AUC | Index | Region name | Hemisphere |
| --- | --- | --- | --- | --- | --- |
| MA | $\Omega_{NR} > \Omega_C$ | 0.970 | 39 | Dorsolateral prefrontal cortex (PFCdl) | Right |
|  |  |  | 41 | Dorsolateral premotor cortex (PMCdl) | Right |
|  |  |  | 46 | Anterior insula (Ia) | Left |
|  |  |  | 82 | Dorsolateral premotor cortex (PMCdl) | Left |
| MA | $\Omega_C > \Omega_{NR}$ | 0.996 | 8 | Orbitolateral prefrontal cortex (PFCol) | Right |
|  |  |  | 15 | Prefrontal polar cortex (PFCpol) | Right |
|  |  |  | 24 | Centrolateral prefrontal cortex (PFCcl) | Right |
|  |  |  | 27 | Posterior cingulate cortex (CCp) | Right |
|  |  |  | 36 | Intraparietal cortex (PCip) | Right |
|  |  |  | 56 | Prefrontal polar cortex (PFCpol) | Left |
|  |  |  | 68 | Posterior cingulate cortex (CCp) | Left |
| DBS | $\Omega_{NR} > \Omega_C$ | 0.724 | 4 | Orbitoinferior prefrontal cortex (PFCoi) | Right |
|  |  |  | 5 | Anterior insula (Ia) | Right |
|  |  |  | 11 | Gustatory cortex (G) | Right |
| DBS | $\Omega_C > \Omega_{NR}$ | 0.974 | 7 | Central temporal cortex (TCc) | Right |
|  |  |  | 9 | Inferior temporal cortex (TCi) | Right |
|  |  |  | 13 | Anterior visual area (ventral, VACv) | Right |
|  |  |  | 19 | Visual area 2 (V2) | Right |
|  |  |  | 21 | Ventral temporal cortex (TCv) | Right |
|  |  |  | 23 | Visual area 1 (V1) | Right |
|  |  |  | 60 | Visual area 2 (V2) | Left |
|  |  |  | 62 | Ventral temporal cortex (TCv) | Left |
|  |  |  | 64 | Visual area 1 (V1) | Left |

**TABLE III.** Mann–Whitney U results for MA dataset (Conscious vs Non-Responsive) and Bonferroni-corrected p-values. Significance levels: * *p*_adj_ < 0.05, ** *p*_ADJ_ < 0.01, *** *p*_ADJ_ < 0.001.

| State | p-value | p-value adj |
| --- | --- | --- |
| $\Omega_{NR} > \Omega_C$ | | |
| 2% Sevo | $5.0 \times 10^{-8}$ | $2.5 \times 10^{-7}***$ |
| 4% Sevo | $8.2 \times 10^{-6}$ | $4.1 \times 10^{-5}***$ |
| Light Ppfl | $3.4 \times 10^{-8}$ | $1.7 \times 10^{-7}***$ |
| Deep Ppfl | $1.1 \times 10^{-8}$ | $5.5 \times 10^{-8}***$ |
| Ketamine | $1.0 \times 10^{-8}$ | $5.1 \times 10^{-8}***$ |
| $\Omega_C > \Omega_{NR}$ | | |
| 2% Sevo | $5.7 \times 10^{-8}$ | $2.9 \times 10^{-7}***$ |
| 4% Sevo | $3.0 \times 10^{-6}$ | $1.5 \times 10^{-5}***$ |
| Light Ppfl | $1.1 \times 10^{-8}$ | $5.3 \times 10^{-8}***$ |
| Deep Ppfl | $4.5 \times 10^{-9}$ | $2.3 \times 10^{-8}***$ |
| Ketamine | $6.9 \times 10^{-9}$ | $3.4 \times 10^{-8}***$ |

**TABLE IV.** Mann–Whitney U results for DBS dataset (Conscious vs Non Responsive) and Bonferroni-corrected p-values. Significance levels: * *p*_ADJ_ < 0.05, ** *p*_ADJ_ < 0.01, *** *p*_ADJ_ < 0.001.

| State | p-value | p-value adj |
| --- | --- | --- |
| $\Omega_{\text{NR}} > \Omega_{\text{C}}$ | | |
| CT high | $2.3 \times 10^{-3}$ | $1.2 \times 10^{-2}$* |
| CT low | $9.8 \times 10^{-3}$ | $4.9 \times 10^{-2}$* |
| VT high | $8.5 \times 10^{-3}$ | $4.2 \times 10^{-2}$* |
| VT low | $4.6 \times 10^{-3}$ | $2.3 \times 10^{-2}$* |
| Off | $4.2 \times 10^{-3}$ | $2.1 \times 10^{-2}$* |
| $\Omega_{\text{C}} > \Omega_{\text{NR}}$ | | |
| CT high | $7.7 \times 10^{-2}$ | $3.9 \times 10^{-1}$ |
| CT low | $8.1 \times 10^{-5}$ | $4.0 \times 10^{-4}$*** |
| VT high | $1.5 \times 10^{-8}$ | $7.5 \times 10^{-8}$*** |
| VT low | $1.7 \times 10^{-8}$ | $8.4 \times 10^{-8}$*** |
| Off | $1.5 \times 10^{-10}$ | $7.4 \times 10^{-10}$*** |

**Fig. 4.**
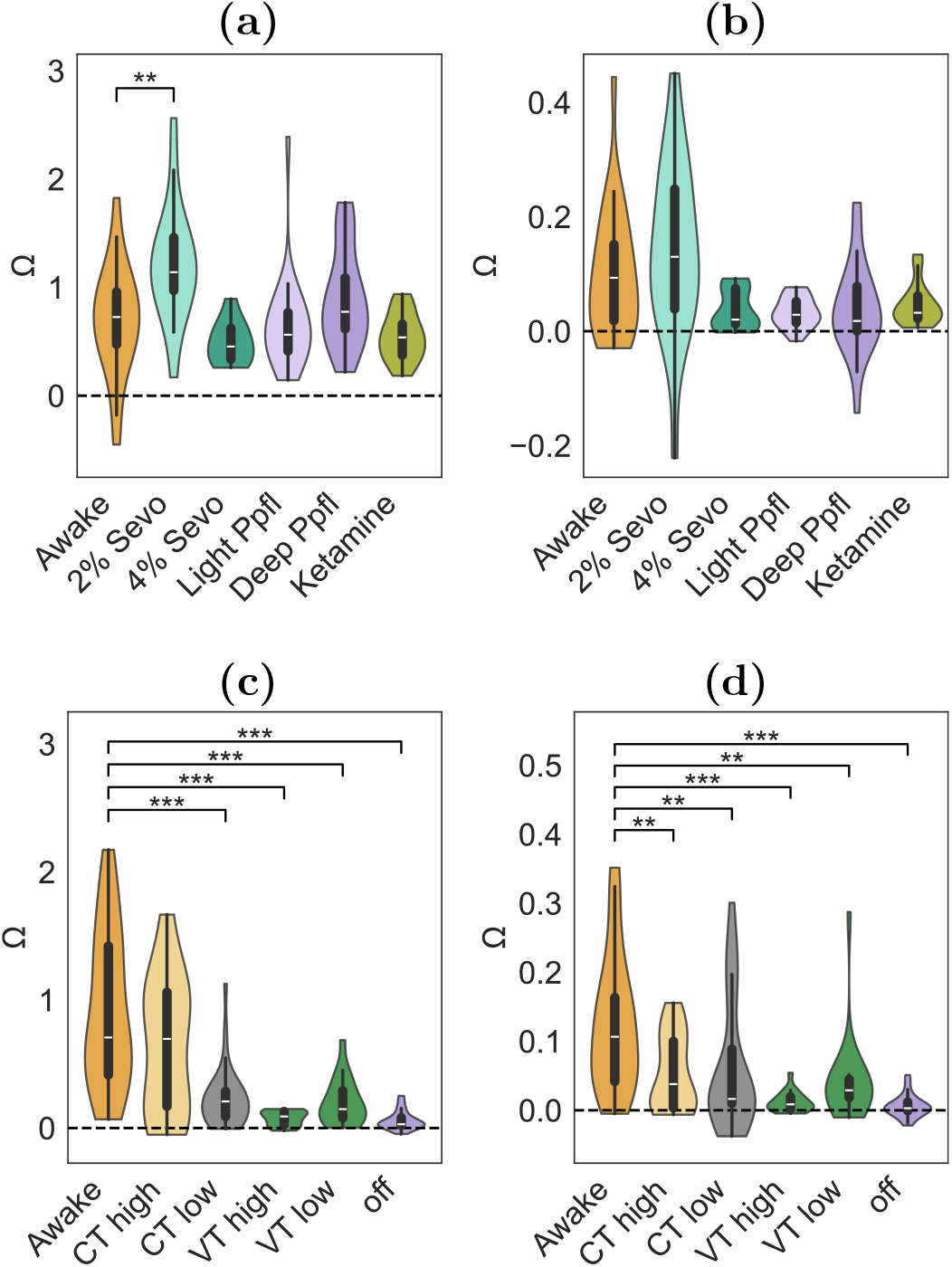
Cross-dataset generalisability of O-information signatures. Violin plots show Ω distributions across scans when optimal *n*-plets identified in one dataset are evaluated on the other. Conscious (C) and non-responsive (NR) macrostates are shown separately for each condition. **(a)** DBS Ω_C_ > Ω_NR_ *n*-plet evaluated on MA scans. **(b)** DBS Ω_NR_ > Ω_C_ *n*-plet evaluated on MA scans. **(c)** MA Ω_C_ > Ω_NR_ *n*-plet evaluated on DBS scans. **(d)** MA Ω_NR_ > Ω_C_ *n*-plet evaluated on DBS scans. PR-AUC values are indicated for each panel. Although the MA Ω_NR_ > Ω_C_ *n*-plet achieves reasonable discrimination on DBS scans (d), the synergy-to-redundancy transition observed in the MA dataset is not preserved: Ω values remain positive across both macrostates, indicating that discrimination arises from a difference in magnitude rather than a sign change.

**Fig. 5.**
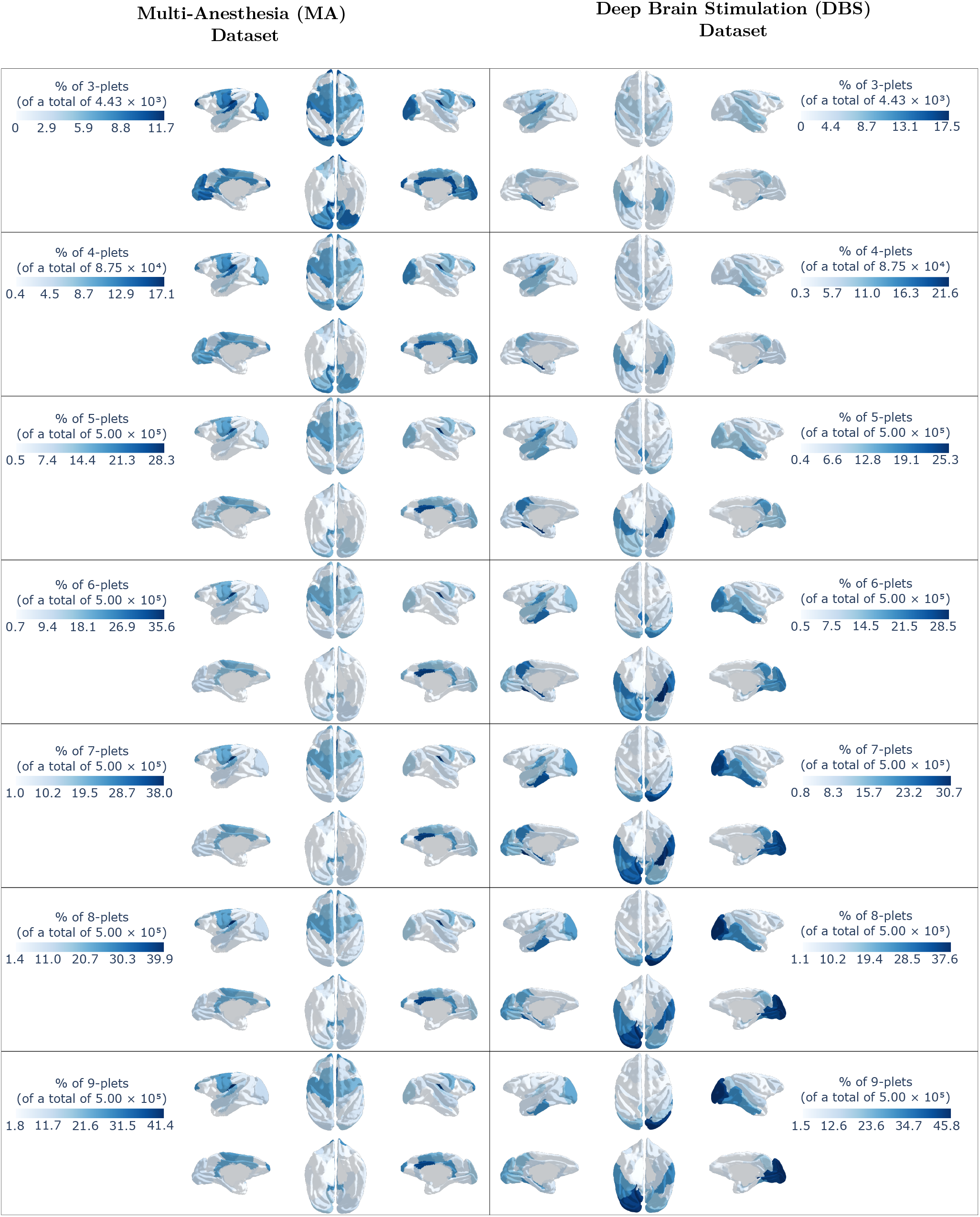
Region participation maps across orders for high-performing Ω_C_ > Ω_NR_ *n*-plets. Brain maps extend the regional participation analysis shown in Fig. 2 to all interaction orders *n* ∈ {3, …, 9} for the Ω_C_ > Ω_NR_ polarity. Columns correspond to the MA (left) and DBS (right) datasets. For each order, regions are coloured by the percentage of filtered, high-performing Ω_C_ > Ω_NR_ *n*-plets in which they appear (see Methods, Section V-E for ranking and sign filtering); the number of *n*-plets contributing to each percentage is indicated in the corresponding panel. Fig. 2 shows the same construction only at the optimal orders for this polarity (MA: *n* = 7; DBS: *n* = 9).

**Fig. 6.**
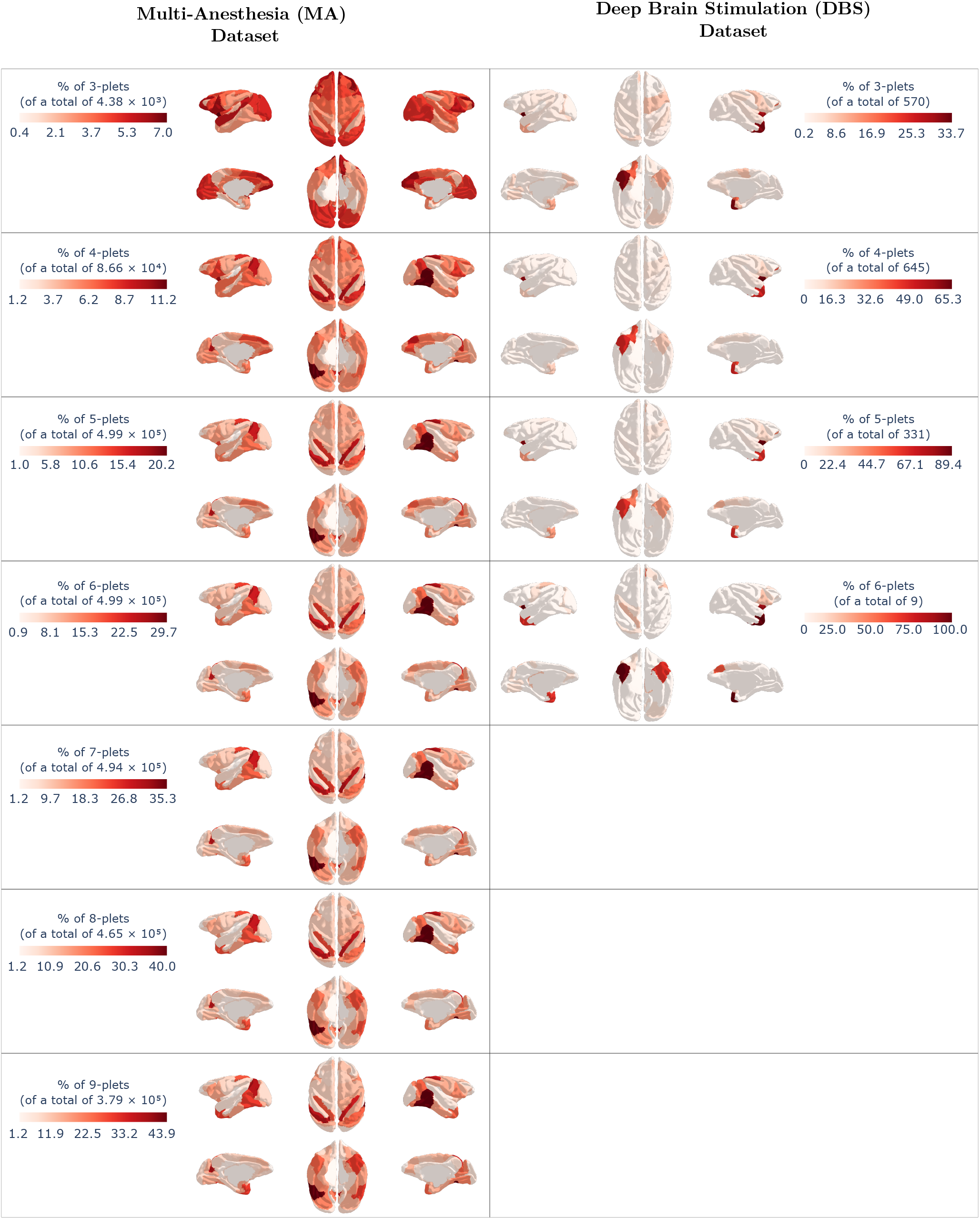
Region participation maps across orders for high-performing Ω_NR_ > Ω_C_ *n*-plets. Same as Fig. 5 but for the Ω_NR_ > Ω_C_ polarity. This figure generalises the Ω_NR_ > Ω_C_ maps shown in Fig. 2 at the corresponding optimal orders (MA: *n* = 4; DBS: *n* = 3). In the DBS dataset, maps are shown only up to *n* = 6 because no candidates satisfy the Ω_NR_ > Ω_C_ sign constraint for *n* ≥ 7 (i.e., no *n*-plets with mean ΔΩ = Ω_NR_ − Ω_C_ > 0 were retained).

**Fig. 7.**
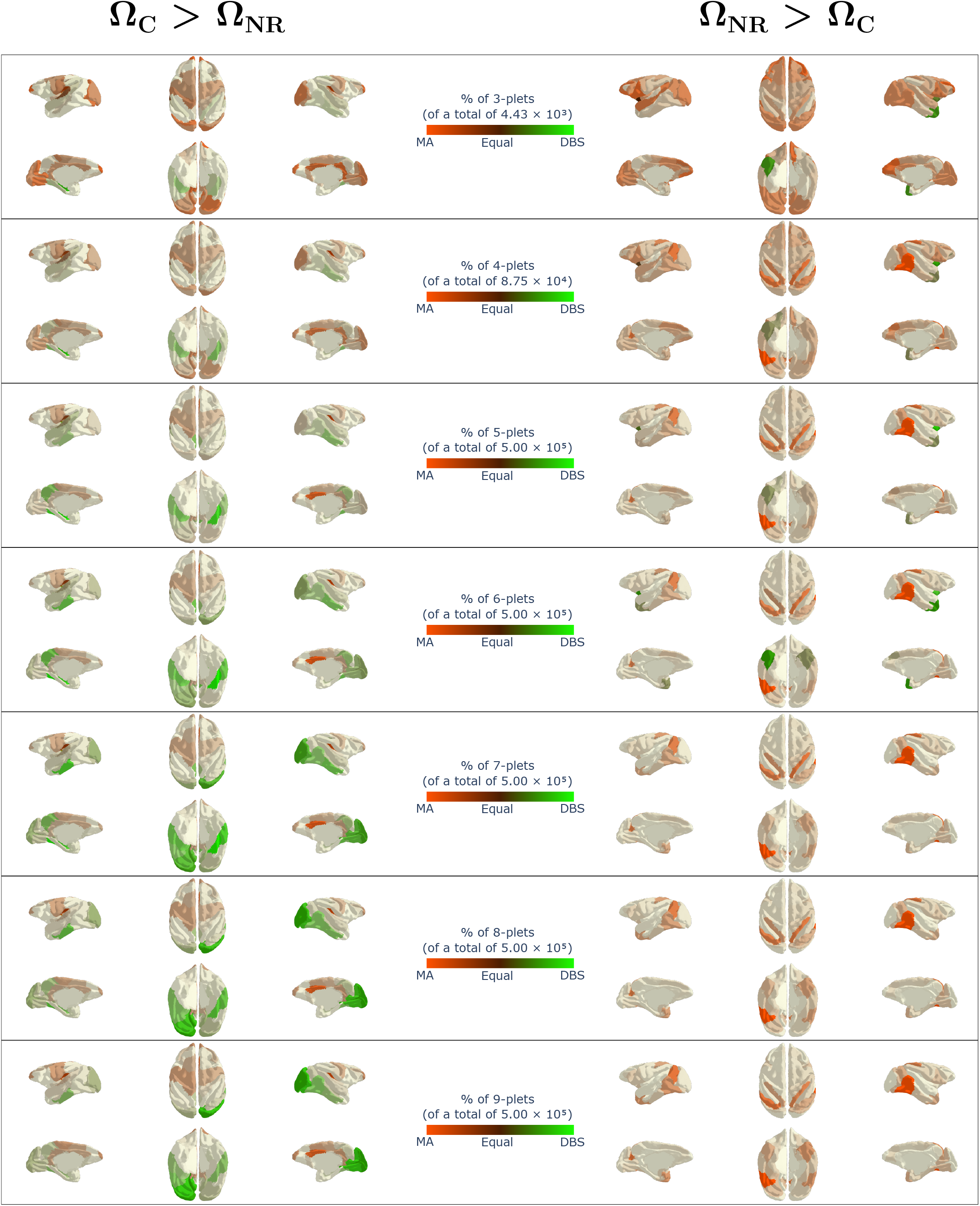
Merged regional participation maps comparing MA- and DBS-dominant higher-order signatures across interaction orders. Brain maps summarise regional participation in high-performing *n*-plets from both datasets across orders *n*∈ *{*3, …, 9}. The left column shows the Ω_C_ > Ω_NR_ polarity and the right column the Ω_NR_ > Ω_C_ polarity. Hue encodes relative dataset dominance, ranging from MA-dominant to DBS-dominant regions, whereas colour intensity reflects overall regional importance. Thus, strongly coloured regions indicate regions that participate frequently in discriminative *n*-plets, with hue indicating whether this participation is more strongly associated with the MA or DBS dataset.

**Fig. 8.**
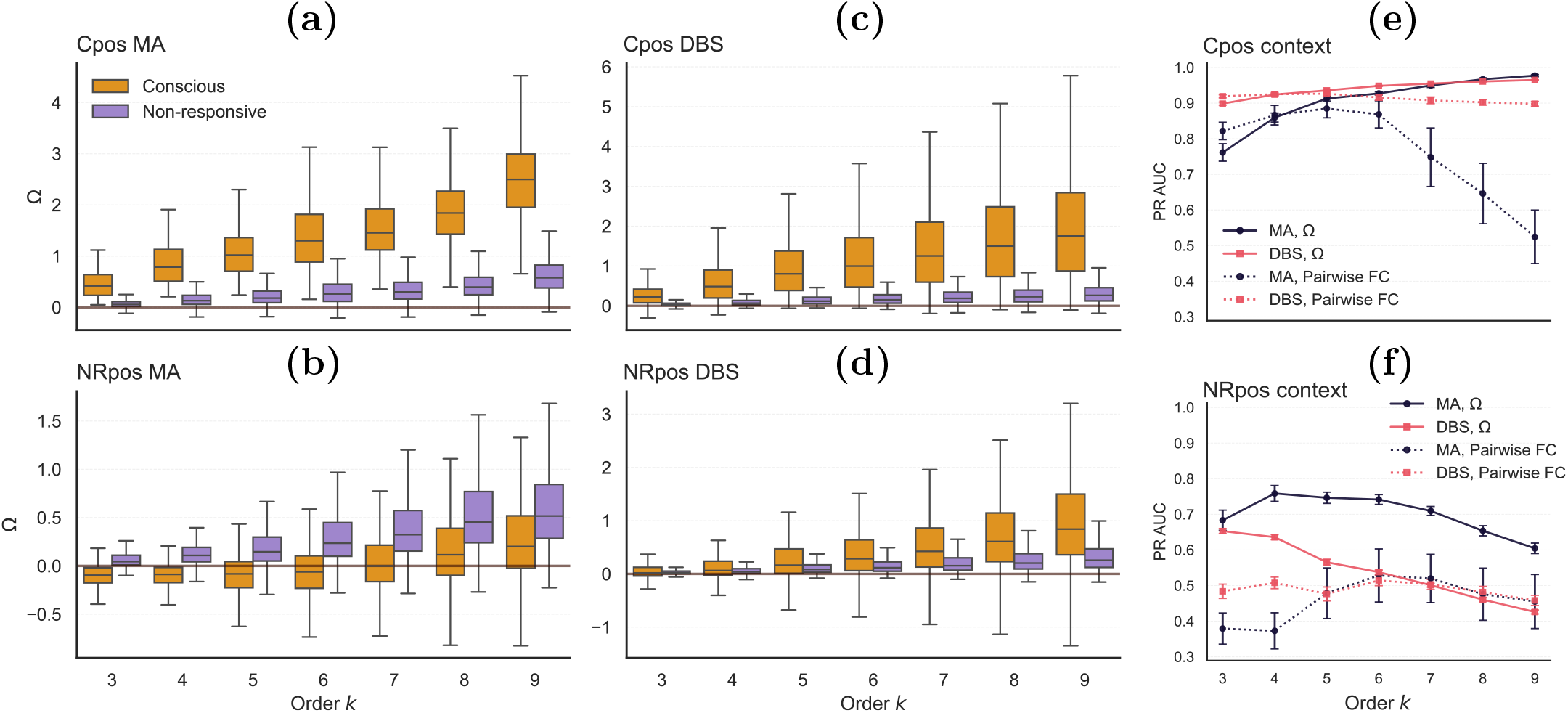
Robustness of order-dependent O-information patterns using top-50 *n*-plets. Unlike Figure 3 in the main text, which shows only the single best *n*-plet per order, this analysis pools the top-50 *n*-plets at each order to assess the robustness of the observed patterns. **(a**,**b)** Distributions of Ω for the top-50 *n*-plets selected under Ω_C_ > Ω_NR_ (a) and Ω_NR_ > Ω_C_ (b) optimisation in the multi-anaesthesia (MA) dataset. For each order *n* (3–9), we selected the top-50 *n*-plets according to PR-AUC (LOPO scheme) and pooled Ω values across all selected *n*-plets and all scans. Boxplots summarise scan-wise values for conscious (C) and non-responsive (NR) scans. **(c**,**d)** Analogous distributions for the deep brain stimulation (DBS) dataset under Ω_C_ > Ω_NR_ (c) and Ω_NR_ > Ω_C_ (d) optimisation. **(e**,**f)** PR-AUC for classifying C versus NR using HOIs (Ω; solid lines) and a pairwise functional connectivity baseline (mean Fisher *z*-values within the same *n*-plet; dashed lines), for Ω_C_ > Ω_NR_ (e) and Ω_NR_ > Ω_C_ (f). The average performance of the top-50 Ω_C_ > Ω_NR_ *n*-plets shows qualitatively similar trends to the single best *n*-plet (Fig. 3e), whereas the top-50 Ω_NR_ > Ω_C_ *n*-plets exhibit more degraded performance compared to the single best *n*-plet (Fig. 3f), suggesting that discriminative Ω_NR_ > Ω_C_ *n*-plets are comparatively rare whilst discriminative Ω_C_ > Ω_NR_ *n*-plets are more abundant.

**Fig. 9.**
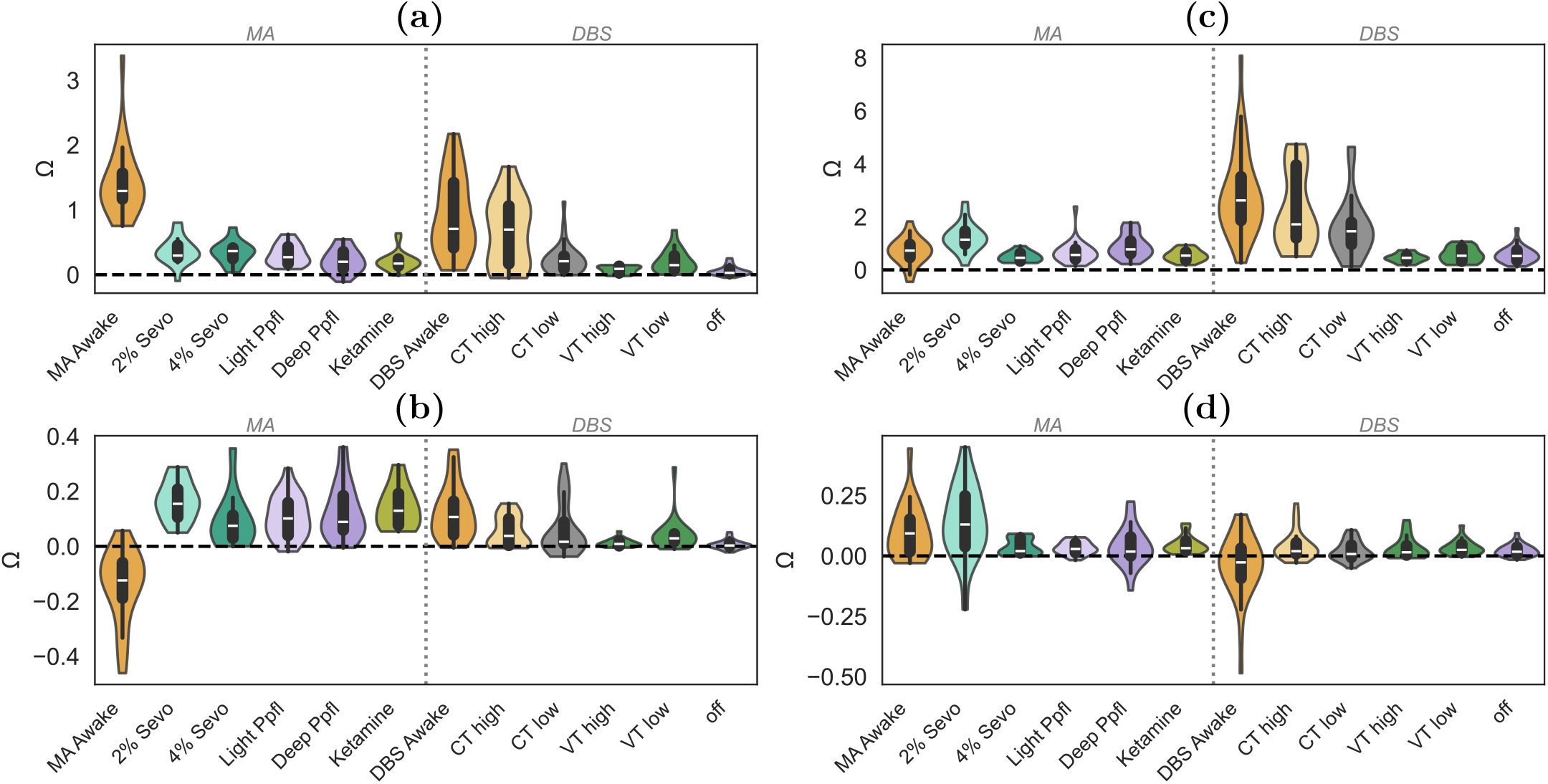
O-information signatures of optimal *n*-plets evaluated on the combined dataset. Violin plots show Ω distributions across all scans from both datasets pooled together (MA+DBS), for optimal *n*-plets identified within each dataset. Conscious (C) and non-responsive (NR) macrostates are shown separately for each condition. **(a)** MA Ω_C_ > Ω_NR_ *n*-plet evaluated on MA+DBS. **(b)** MA Ω_NR_ > Ω_C_ *n*-plet evaluated on MA+DBS. **(c)** DBS Ω_C_ > Ω_NR_ *n*-plet evaluated on MA+DBS. **(d)** DBS Ω_NR_ > Ω_C_ *n*-plet evaluated on MA+DBS. PR-AUC values are indicated for each panel.

The results reveal a strong asymmetry in cross-dataset transferability. *n*-plets identified in the MA dataset generalise well to the DBS dataset: the MA Ω_C_ > Ω_NR_ *n*-plet achieves PR-AUC = 0.915 on DBS scans, and the MA Ω_NR_ > Ω_C_ *n*-plet reaches PR-AUC = 0.832. In contrast, *n*-plets identified in the DBS dataset fail to discriminate states in the MA dataset: the DBS Ω_C_ > Ω_NR_ *n*-plet attains only PR-AUC = 0.212 on MA scans, and the DBS Ω_NR_ > Ω_C_ *n*-plet reaches PR-AUC = 0.312.

Generalisability is also strongly polarity-dependent, and crucially, the violin plot distributions in Fig. 4 reveal that numerical transfer and mechanistic transfer can dissociate. For the Ω_C_ > Ω_NR_ polarity, both numerical performance and the underlying informational pattern transfer robustly: across all cross-dataset evaluations, conscious scans show higher positive Ω than non-responsive scans (Fig. 4a,c), consistent with the within-dataset signature of elevated redundancy in wakefulness. When both datasets are combined, the MA Ω_C_ > Ω_NR_ *n*-plet achieves PR-AUC = 0.873 and the DBS *n*-plet reaches PR-AUC = 0.801, confirming that this redundancy-based signature reflects a conserved feature of conscious states across diverse perturbations.

By contrast, the Ω_NR_ > Ω_C_ polarity shows a dissociation between numerical and mechanistic generalisability. Although the MA Ω_NR_ > Ω_C_ *n*-plet achieves reasonable discrimination on DBS scans (PR-AUC = 0.832), the synergy-to-redundancy transition that defines this signature in the MA dataset (conscious: negative Ω, non-responsive: positive Ω) is not preserved: when evaluated on DBS scans, Ω values remain positive across both macrostates, and discrimination arises from a difference in magnitude rather than a sign change (Fig. 4d). This pattern holds across all four cross-dataset panels for the Ω_NR_ > Ω_C_ polarity (Fig. 4b,d): none show a negative-to-positive Ω transition, which seems to indicate that the synergy-to-redundancy shift is specific to the MA context and does not constitute a generalisable feature of consciousness loss. Accordingly, Ω_NR_ > Ω_C_ *n*-plets collapse when datasets are combined (MA: PR-AUC = 0.313; DBS: PR-AUC = 0.351), consistent with the within-dataset weakness of this polarity in DBS (PR-AUC = 0.724) and the considerably fewer candidate *n*-plets satisfying the target contrast in that dataset (570 vs. thousands in other contexts).

## III. Discussion

In this study, we demonstrate that HOIs, quantified through O-information, provide a robust and generalizable signature of brain states associated with states of consciousness. Across two independent datasets (anaesthesia with different molecular agents and thalamic deep brain stimulation), we show that multiregional informational interactions reliably classify different states of consciousness, outperforming conventional pairwise functional connectivity measures. This is not driven by a single canonical pattern, but rather by distinct yet complementary re-configurations of the brain’s informational architecture, involving both redundancy- and synergy-dominated regimes. Our findings are consistent with prior literature, which indicates that HOIs within brain networks may play a crucial role in elucidating alterations in brain dynamics across varying states of consciousness and cognitive states [16], [38], [31], [35].

We identified two consistent ways in which HOIs differentiate wakefulness from non-responsiveness. First, under Ω_NR_ > Ω_C_ optimisation, discriminative *n*-plets exhibit a transition from synergy-dominated interactions (Ω < 0) in awake scans to redundancy-dominated interactions (Ω > 0) under anaesthesia. This synergy-to-redundancy reconfiguration has been reported previously [35], and can be related to the reduced emergent character of neural dynamics in patients with disorders of consciousness [39] and the breakdown of information integration under general anaesthesia across mammalian brains [12]. Second, Ω_C_ > Ω_NR_ *n*-plets show elevated redundancy in wakefulness that decreases, but remains positive (Ω > 0), in non-responsive scans. This pattern was the most robust across datasets and achieved near-perfect discrimination, particularly at high interaction orders. Across datasets, Ω_C_ > Ω_NR_ *n*-plets yielded the most reliable macrostate discrimination, whereas Ω_NR_ > Ω_C_ *n*-plets showed less stability and context-dependent performance.

Regional participation patterns further distinguished the two optimisation polarities and revealed an apparent paradox: despite highly consistent Ω distributions across datasets, the anatomical distributions of high-performing *n*-plets differed markedly. In the MA dataset, Ω_C_ > Ω_NR_ *n*-plets showed concentrated involvement of frontal and cingulate cortices (peak frequency 37.9%), whilst Ω_NR_ > Ω_C_ *n*-plets exhibited widespread participation across cortex with stronger involvement in visual and motor regions (peak frequency 11.2%). In the DBS dataset, Ω_C_ > Ω_NR_ *n*-plets were dominated by visual cortex (45.8%), whilst Ω_NR_ > Ω_C_ *n*-plets were more spatially concentrated in right temporal cortex (33.7%).

The regional concentration also differed between polarities: Ω_C_ > Ω_NR_ *n*-plets, despite being larger (*n* = 7-9), showed higher peak regional frequencies, indicating that certain regions are repeatedly selected, whilst Ω_NR_ > Ω_C_ *n*-plets showed more distributed participation (MA) or failed to achieve robust discrimination (DBS).

The dependence of discrimination performance on interaction order differed markedly between optimization polarities. For Ω_C_ > Ω_NR_ *n*-plets, separability improved monotonically with order, reaching near-perfect classification at *n* = 7 (MA, PR-AUC = 0.996) and *n* = 9 (DBS, PR-AUC = 0.974). The pairwise functional connectivity baseline showed a different behaviour: in the MA dataset, pairwise features performed comparably to Ω at low orders but then declined dramatically at higher orders, whilst in the DBS dataset pairwise features remained relatively stable but failed to improve with increasing order as Ω-based features did. This suggests that at low orders, redundancy may be partially captured by strong pairwise correlations, potentially reflecting anatomical connectivity patterns. However, as subset size increases, the multivariate redundancy structure requires higher-order measures to detect, and pairwise descriptions become increasingly insufficient.

In contrast, Ω_NR_ > Ω_C_ *n*-plets peaked at low orders (*n* = 3-4) and declined progressively as more regions were added. The boxplot distributions in Figure 3c,d show that even the best-performing *n*-plets at each order exhibit this pattern, with Ω values becoming positive in both macrostates and distributions increasingly overlapping from *n* ≥ 5 onwards in MA. This indicates that the synergy-to-redundancy transition dissolves at higher orders even among optimally selected regional subsets, suggesting this signature is intrinsically a low-order phenomenon. When additional regions are included, the emergent character may be obscured if those regions do not contribute coherently to the synergistic interaction. For Ω_NR_ > Ω_C_ *n*-plets, pairwise features were consistently and substantially worse than Ω across all orders in both datasets, demonstrating that synergistic interactions are fundamentally invisible to pairwise correlations.

The cross-dataset generalisation analysis revealed an important distinction between generalisability of the studied higher-order signatures. MA-derived *n*-plets generalise robustly to DBS (PR-AUC ≥ 0.83 for both polarities), whilst DBS-derived *n*-plets fail to discriminate states in MA (PR-AUC ≤ 0.31). This asymmetry likely reflects the broader variance captured by the MA dataset, which includes three anesthetic agents with fundamentally different molecular mechanisms (ketamine, propofol, and sevoflurane), all converging on behavioural unresponsiveness. The DBS dataset, by contrast, examines only propofol anaesthesia modulated by thalamic stimulation. The MA optimization procedure thus samples a wider space of non-responsive states, enabling identification of *n*-plets whose Ω signatures capture more generalizable features of loss of consciousness.

The Ω_C_ > Ω_NR_ signature generalises both numerically and mechanistically: across all cross-dataset evaluations, conscious scans show higher positive Ω than non-responsive scans, and discrimination performance remains high when datasets are combined (MA: PR-AUC = 0.873; DBS: PR-AUC = 0.801). This confirms that elevated redundancy in conscious states reflects a conserved informational property across diverse perturbations. Notably, this signature places high-voltage central thalamus stimulation during propofol anaesthesia closer to wakefulness than to anesthetized conditions, consistent with the behavioural arousal effects reported in the original DBS study [20].

By contrast, the Ω_NR_ > Ω_C_ signature shows a dissociation between numerical and mechanistic generalisability. Although the MA *n*-plet achieves PR-AUC = 0.832 on DBS scans, the synergy-to-redundancy transition that defines this signature in MA (conscious: negative Ω, non-responsive: positive Ω) is absent when evaluated on DBS data: Ω values remain positive in both macrostates, and discrimination reflects a magnitude difference rather than a sign change. No cross-dataset evaluation for this polarity recovers the sign transition, confirming that the synergy-to-redundancy shift is specific to the MA context rather than a general feature of consciousness loss. Accordingly, Ω_NR_ > Ω_C_ *n*-plets collapse when datasets are combined (MA: PR-AUC = 0.313; DBS: PR-AUC = 0.351).

Several limitations should be considered. First, our analyses rely on group-level optimization and evaluation, which may obscure individual variability in higher-order interaction patterns. Second, our analyses rely on Gaussian copula-based estimators (THOI) [35], which may not capture dependencies in higher moments or non-Gaussian distributions. However, Gaussianity is a reasonable first approximation for fMRI data [40], and alternative estimators (e.g. kernel-based, non-parametric) implemented in JIDT [41] or the HOI toolbox [37] could provide more robust HOI estimations in future work. Third, preprocessing differences between datasets may contribute to observed differences in Ω signatures. The MA dataset employed global signal regression, whilst the DBS dataset did not, which may influence the balance between synergistic and redundant interactions and could partially explain the greater prevalence of synergy-dominated *n*-plets in MA. Future studies with harmonized preprocessing pipelines would help isolate genuine biological differences from methodological artifacts. Fourth, whilst O-information provides a useful summary of redundancy–synergy balance, it does not fully decompose the underlying informational structure, and future work using more refined decompositions may yield additional insights. Finally, our study focuses on resting-state fMRI in non-human primates, and the extent to which these findings generalize to humans and task-based paradigms remains to be established, but recent findings in deep anaesthesia suggest that this could be the case [35].

## IV. Conclusion

In summary, we show that higher-order informational interactions provide a powerful and generalizable signature of consciousness, capturing aspects of brain dynamics that remain inaccessible to pairwise analyses. Two complementary informational reconfigurations distinguish conscious from non-responsive states: a conserved elevation of redundancy in wakefulness that generalizes across anesthetic agents and thalamic neuromodulation, and a context-dependent synergy-to-redundancy transition prominent under certain pharmacological manipulations. These findings align with a growing body of work highlighting the relevance of higher-order dependencies in complex systems, including recent advances in information-theoretic and topological characterizations of multivariate interactions [8], [42], [43], [44].

Our results support the view that consciousness is not merely reflected in changes in specific brain regions or pairwise functional connectivity, but in the reconfiguration of higher-order interaction structures that govern collective neural dynamics. Within this framework, measures such as the O-information and related descriptors provide principled tools to disentangle synergistic and redundant contributions across scales, offering a unifying perspective on the organization of brain activity.

Future work should extend this approach along several key directions. First, incorporating subcortical regions, in particular the thalamus (which has been shown to be central in the DBS experiment [20], [45]), and integrating multimodal datasets (including fMRI, electrophysiology, and receptor density maps) will be critical to achieving a more comprehensive account of brain organization. Second, embedding these empirical findings within mechanistic whole-brain models will allow us to link anatomical structure, dynamical regimes, and neuromodulatory influences to the emergence and modulation of conscious states [46], [47], [48], [49]. Third, exploring structural connectivity–functional connectivity coupling, particularly at low interaction orders where pairwise and higher-order features converge, may reveal whether anatomical constraints shape the observed informational architecture. Finally, it remains essential to carefully distinguish between observed statistical dependencies and their underlying generative mechanisms, as higher-order statistical structure can emerge from lower-order interactions in complex systems [50]. Bridging this gap will be crucial for advancing from statistical to mechanistic accounts of consciousness.

## V. Methods

### A. Datasets

We analyzed two previously reported datasets [22], [20].

#### 1) Multi-Anaesthesia Dataset

##### a) Animals

Five adult rhesus macaques (*Macaca mulatta*; one male, four females; 5–8kg; 8–12yr) were included across six arousal conditions: awake, ketamine, light propofol, deep propofol, light sevoflurane, and deep sevoflurane anaesthesia. Three monkeys were used per condition: awake (monkeys A, K, J), ketamine (K, R, Ki), propofol (K, R, J), and sevoflurane (Ki, R, J). Each monkey underwent resting-state fMRI acquisitions on separate days; some monkeys contributed data to multiple conditions. Sex was not considered due to limited sample size. All procedures complied with the European Directive 2010/63/EU and NIH Guide for the Care and Use of Laboratory Animals, and were approved by the institutional Ethical Committee (CETEA #10-003 and #12-086; Commissariat à l’Énergie Atomique et aux Énergies Alternatives, Fontenay-aux-Roses, France). Acquisition details are available in the original publications [21], [22].

##### b) Anaesthesia Protocol

Monkeys received ketamine, propofol, or sevoflurane anaesthesia [21], [22], with two levels for propofol and sevoflurane (light and deep). Anaesthesia depth was determined using the monkey sedation scale, based on spontaneous movements, response to stimuli (presentation, shaking/prodding, toe pinch), and corneal reflex [51]. Clinical scores were assessed at session onset and conclusion, alongside continuous EEG monitoring. Monkeys were intubated, ventilated, and physiological parameters (heart rate, noninvasive blood pressure, oxygen saturation, respiratory rate, end-tidal CO_2_, and cutaneous temperature) were continuously monitored (Maglife, Schiller, France).

Ketamine, deep propofol, and deep sevoflurane anaesthesia induced unresponsiveness to stimuli, consistent with general anaesthesia. Ketamine was administered intramuscularly (20mg/kg) for induction, followed by continuous intravenous infusion (15–16mg·kg^−1^·h^−1^). Atropine (0.02mg/kg, i.m.) was administered 10min prior to reduce secretions. Propofol was delivered as an intravenous bolus (5–7.5mg/kg) followed by target-controlled infusion (TCI; light: 3.7–4.0µg/ml, deep: 5.6–7.2µg/ml) using the Paedfusor pharmacokinetic model. Sevoflurane anaesthesia was induced with ketamine (20mg/kg, i.m.) followed by inhalation (light: 2.2/2.1% insp/exp; deep: 4.4/4.0% insp/exp). Scanning commenced 80min post-induction to allow ketamine washout. Muscle relaxation was achieved with cisatracurium (0.15mg/kg bolus, 0.18mg·kg^−1^·h^−1^ infusion) during ketamine and light propofol sessions to minimize motion artifacts.

##### c) Functional MRI Acquisition

Monkeys were trained to sit in sphinx position in a primate chair. Awake sessions involved eye monitoring at 120Hz (Iscan Inc., USA) to ensure wakefulness; eye movements were not regressed out. During anaesthesia, animals were mechanically ventilated with physiological monitoring. Prior to scanning, MION contrast agent (10 mg/kg, i.v.) was administered. Imaging was performed on a 3-T Siemens Tim Trio scanner using a custom transmit-receive surface coil. Gradient-echo EPI parameters: TR=2400ms, TE=20ms, voxel size = 1.5 mm^3^, 500 volumes per run.

##### d) Data Preprocessing and Time Series Extraction

A total of 157 resting-state fMRI runs were acquired [22]. Preprocessing included reorientation, realignment, and rigid coregistration to the monkey MNI template using Python and FSL. Global signal regression was applied. After expert visual quality control, 119 runs were retained: awake (24), ketamine (22), light propofol (21), deep propofol (23), light sevoflurane (18), deep sevoflurane (11). Data were parcellated using the Regional Map parcellation (82 cortical ROIs, 41 per hemisphere) [52]. Voxel time series were band-pass filtered (0.0025–0.05Hz) and notch filtered at 0.03Hz. Expert quality control included inspection of time series, static functional connectivity, and Fourier spectra, rejecting trials with stereotyped oscillations or residual motion artifacts.

#### 2) DBS Dataset

##### a) Animals

Five male rhesus macaques (9–17yr, 7.5–9.1kg) were included: three for awake (non-DBS) experiments (B, J, Y) and two for DBS experiments (N, T). Only males were included to control for hormonal fluctuations. Procedures adhered to EU directive 2010/63/UE and related protocols [20].

##### b) Anaesthesia Protocol

Anaesthesia was induced with ketamine (10mg/kg, i.m.) and dexmedetomidine (20µg/kg, i.m.), followed by propofol TCI (Monkey T: 4.6–4.8µg/ml; Monkey N: 4.0–4.2µg/ml) [20]. Awake fMRI data were obtained from three additional monkeys.

##### c) Deep Brain Stimulation Protocol

Two macaques (N, T) were implanted with clinical DBS electrodes (Medtronic 3389; 1.5-mm contacts, 0.5-mm spacing, 1.27-mm diameter) targeting the right centromedian thalamus using stereotactic guidance (BrainSight, Rogue, Canada) based on macaque atlases [53], [54] and preoperative/intraoperative MRI (MPRAGE, TR=2200ms, TI=900ms, 0.80-mm isotropic). Electrode location was verified via in vivo MRI and postmortem in one animal. DBS-fMRI experiments commenced ≥20 days post-implantation. Stimulation was delivered via external stimulator (DS8000, WPI) with fixed parameters: frequency 130.208Hz, monopolar waveform, pulse width 320*µ*s (N) or 140*µ*s (T). Two voltage regimes were applied: low (3V) and high (5V), targeting either centromedian (CT) or ventrolateral (VT) thalamic nuclei under anaesthesia. High CT DBS significantly increased heart rate and blood pressure. Imaging was also acquired during wakefulness and anaesthesia without stimulation (“off” condition).

##### d) Functional MRI Acquisition

Monkeys were trained to sit in sphinx position in a primate chair. Awake sessions involved eye monitoring at 120Hz (Iscan Inc., USA) to ensure wakefulness; eye movements were not regressed out. During anaesthesia, animals were mechanically ventilated with physiological monitoring. Scanning used a 3-T Siemens Prisma Fit with an 8-channel phased-array surface coil. EPI parameters: TR=1250ms, TE=14.2ms, voxel size=1.25mm isotropic, 500 volumes per run. EEG was acquired with 13-channel MR-compatible caps but not analyzed here [20].

##### e) Data Preprocessing and Time Series Extraction

A total of 199 runs were acquired [20]. Preprocessing employed Pypreclin [55]: slice-timing and B0 correction, reorientation, realignment, resampling (1mm isotropic), masking, coregistration to MNI macaque template, and 3-mm Gaussian smoothing. Quality control resulted in 156 runs for analysis: awake (36), DBS-off (28), low CT DBS (31), low VT DBS (18), high CT DBS (25), high VT DBS (18). Data were parcellated using the Regional Map parcellation (82 cortical ROIs, 41 per hemisphere) [52]. Voxel time series were band-pass filtered (0.0025–0.05Hz) and notch filtered at 0.03Hz. Expert quality control included inspection of time series, static functional connectivity, and Fourier spectra, rejecting trials with stereotyped oscillations or residual motion artifacts.

#### B. Multivariate Information Theory and Higher-Order Interactions

Information theory provides a rigorous framework to quantify statistical dependencies within multivariate systems, extending well beyond the pairwise domain. At its foundation lies Shannon’s differential entropy, which for a continuous random variable *X* with density *p*(*x*) is defined as

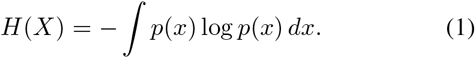

For a multivariate random vector **X**_*n*_ = (*X*_1_, …, *X*_*n*_), *H*(**X**_*n*_) denotes the joint entropy, while *H*(*X*_*j*_) refers to the marginal entropy of the *j*-th component.

Pairwise dependencies are captured by the mutual information between two variables, *X* and *Y*, which admits the following equivalent expressions:

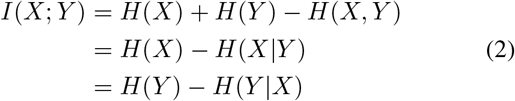

In multivariate systems (i.e., *n* > 2), the expressions in (2) give rise to complementary higher-order indicators.

#### 1) Total Correlation and Dual Total Correlation

Two classical extensions of mutual information are the *total correlation* (TC) [56] and the *dual total correlation* (DTC) [57], defined respectively as:

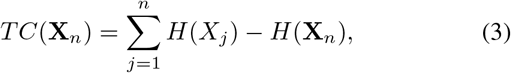

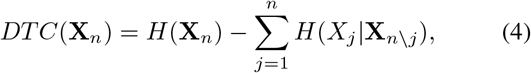

where **X**_*n\j*_ denotes the full vector excluding variable *X*_*j*_. Both are non-negative measures: TC quantifies the extent to which the system (i.e., **X**_*n*_) is collectively constrained, whereas DTC characterizes the amount of shared randomness among the components of **X**_*n*_.

#### 2) O-information

Building on TC and DTC, Rosas et al. [9] introduced the O-information (Ω), a metric specifically designed to capture the balance between redundancy and synergy in high-dimensional systems:

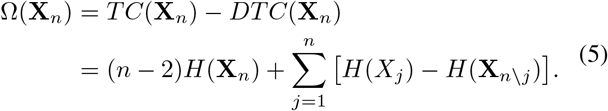

If Ω(**X**_*n*_) > 0, redundancy dominates the system, indicating that shared information is prevalent across variables. Conversely, if Ω(**X**_*n*_) < 0 the system is dominated by synergistic dependencies in the sense that collective information emerges only when variables are considered jointly.

#### 3) Practical Computation with THOI

Despite their conceptual clarity, computing these higher-order metrics is notoriously challenging due to the exponential scaling of possible *n*-variable subsets and the difficulties of estimating high-dimensional entropies. To overcome these challenges, we rely on the recently introduced THOI (Torch High-Order Interactions) library [35], which leverages Gaussian copula entropy estimators [58] and highly optimized batch-processing routines on modern hardware (CPU/GPU/TPU).

The Gaussian copula estimator avoids direct density estimation by transforming each variable to a standard normal via its empirical cumulative distribution function, preserving rank dependencies while discarding marginal structure. Under this transformation, the joint entropy can be approximated using the covariance matrix ∑ of the transformed variables as:

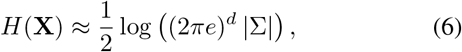

where *d* is the dimensionality of the system. This approach provides a scalable and robust estimate of multivariate entropy that is well-suited for high-dimensional settings.

`THOI` enables the efficient estimation of *TC, DTC*, and Ω across a large number of subsets of variables, thereby making exhaustive or heuristic analyses of higher-order interactions feasible even in moderately high-dimensional systems. In this work, all higher-order informational quantities reported were computed using `THOI`.

#### C. Pairwise Baseline

As a pairwise functional connectivity (FC) baseline, we computed Pearson correlations between all pairs of regions studied and applied the Fisher *z*-transform

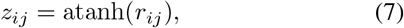

where *r*_*ij*_ denotes the Pearson correlation between regions *i* and *j*. Fisher-*z* transformed correlations are widely used to characterize large-scale brain networks in fMRI [59], [60], [61], [62]. In this work, these pairwise connectivity estimates were summarized at the *n*-plet level (e.g., as mean within-*n*-plet connectivity) and used as a canonical FC benchmark. All comparative analyses contrast this Fisher-*z*-based pairwise functional connectivity against Ω.

#### D. n-plet Selection and Optimization

Due to the combinatorial explosion in the number of possible sets of brain regions, and to maintain interpretability, we restricted our search to *n*-plets with sizes *n* = 3, …, 9. Candidate *n*-plets were identified using a heuristic search based on simulated annealing [10], as implemented in THOI and adapted to our setting. Simulated annealing is a stochastic local search procedure that iteratively modifies a current candidate solution and accepts changes that improve the objective, while occasionally accepting worse solutions with a probability that decreases over time (the “cooling schedule”), thereby reducing the risk of getting trapped in poor local optima.

The search scheme was organized around pairs of scans drawn from the two macrostates: C and NR. For each subject pair and each *n* ∈ {3, …, 9}, we used simulated annealing to identify an *n*-plet of regions that maximized the difference in O-information between the two scans. We considered two optimization polarities:

- Ω_C_ > Ω_NR_, in which higher Ω values are associated with the conscious state, and
- Ω_NR_ > Ω_C_, in which higher Ω values are associated with the non-responsive state.

For each subject pair, *n*-plet size, and polarity, simulated annealing was run to approximate the *n*-plet that maximized the signed difference in O-information between the two scans. The annealing procedure operated for 10,000 steps per run, was repeated 200 times to reduce stochastic variability, and employed an early-stopping threshold of 1,000 steps without improvement. These parameters were selected to ensure convergence whilst remaining computationally feasible.

The resulting collection of selected *n*-plets was then evaluated in a leave-one-pair-out (LOPO) manner. For each candidate *n*-plet, we computed Ω for all scans in the corresponding dataset, excluding (i) the specific C-NR scan pair used to discover that *n*-plet and (ii) the intermediate three-volt stimulation condition in the DBS dataset. This yielded, for each scan, a single scalar value Ω associated with that *n*-plet, which we treated as the scan-wise score used to classify C and NR states.

##### 1) n-plet evaluation

To quantify how well a given *n*-plet separates C from NR, we used the area under the precision-recall curve (PR-AUC). Unlike the receiver operating characteristic (ROC) curve, which plots true positive rate against false positive rate, the precision-recall curve plots precision (fraction of positive predictions that are correct) against recall (fraction of true positives detected). For a chosen polarity (Ω_C_ > Ω_NR_ or Ω_NR_ > Ω_C_), one macrostate is treated as the positive class and the other as the negative class. Varying a decision threshold on Ω produces different trade-offs between avoiding false alarms (precision) and correctly detecting the positive macrostate (recall). The area under the PR curve provides a single score between 0 and 1, where PR-AUC = 1 indicates perfect separation between macrostates. We use PR-AUC as our summary metric because it directly captures how well an *n*-plet, evaluated through Ω, distinguishes one macrostate from the other and remains informative when the numbers of C and NR scans are imbalanced.

Because the simulated annealing procedure is stochastic, the same *n*-plet could be rediscovered across different subject pairs. For each unique *n*-plet, we therefore averaged its PR-AUC score across all discovery pairs, and retained a single non-duplicated entry per *n*-plet. Within each dataset and polarity, we then selected the *n*-plet with the highest mean PR-AUC as the optimal *n*-plet for that configuration.

#### E. Region Importance Maps

To identify which regions are consistently selected in discriminative higher-order interactions, we constructed regional importance maps for the same *n*-plet sizes as the four optimal configurations (*n* = 3, 4, 7, and 9). These maps summarize, for each region, how frequently it participates in *n*-plets that reliably distinguish C from NR states based on O-information.

We obtained these maps using the following sampling and evaluation procedure:

- For *n* = 3 and *n* = 4, we retained the top 5% of *n*-plets ranked by PR-AUC (approximately 4,400 and 87,500 n-plets, respectively). For *n* = 7 and *n* = 9, exhaustive evaluation of the top 5% is computationally infeasible, so we retained the top 5 × 10^5^ *n*-plets (corresponding to 1% of the sampled combinations). For each *n*-plet and dataset, we computed Ω for all scans (excluding the intermediate 3V condition in DBS) and quantified its discriminability between C and NR using PR-AUC, following the same polarity conventions as in the optimisation stage. That is, for sizes *n* = 3 (DBS) and *n* = 4 (MA), which correspond to Ω_NR_ > Ω_C_ optimal *n*-plets, larger Ω values were interpreted as evidence for the non-responsive state; conversely, for sizes *n* = 7 (MA) and *n* = 9 (DBS), which correspond to Ω_C_ > Ω_NR_ optimal *n*-plets, larger Ω values were interpreted as evidence for the conscious state.
- For *n* = 3 and *n* = 4, we retained the top 5% of *n*-plets ranked by PR-AUC. For *n* = 7 and *n* = 9, we retained the 5 × 10^5^ *n*-plets with the highest PR-AUC. Within each such tail of high-performing *n*-plets, we further enforced consistency with the expected polarity by filtering according to the signed difference

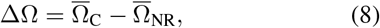

where the overline denotes averaging across scans within each macrostate. For the Ω_NR_ > Ω_C_ polarity (sizes *n* = 3 and *n* = 4), we only retained *n*-plets with ΔΩ < 0; for the Ω_C_ > Ω_NR_ polarity (sizes *n* = 7 and *n* = 9), we only retained *n*-plets with ΔΩ > 0.
- Finally, for each dataset, polarity, and *n*-plet size, we computed a regional importance score by counting how many times each region appeared across all *n*-plets in the filtered tail and dividing by the total number of *n*-plets in that tail:

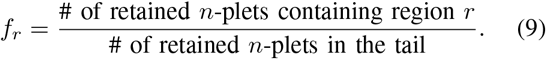

We interpret *f*_*r*_ as the fraction of high-performing *n*-plets in which region *r* participates. For visualization, these values were expressed as percentages (100 × *f*_*r*_) in Fig. 2.

### F. Order-dependent separability analysis

To assess how state separability, measured by PR-AUC, varied with *n*-plet order, we focused on the best-performing *n*-plets identified in the optimisation stage (Section V-D). For each order *n* ∈ {3, …, 9}, dataset (MA, DBS), and polarity (Ω_C_ > Ω_NR_, Ω_NR_ > Ω_C_), we selected the *n*-plet with the highest mean PR-AUC. For each of these *n*-plets, we then computed Ω for all scans in the corresponding dataset (excluding the intermediate 3V condition in DBS as described above). This yielded, for each scan and order *n*, a set of Ω values associated with the top-performing *n*-plet. These values were used to summarise the distribution of Ω across scans and orders in Fig. 3a–d.

In parallel, we quantified classification performance for the same set of *n*-plets. For each order, we evaluated how well its Ω values discriminated conscious (C) from non-responsive (NR) scans using PR-AUC, treating the appropriate macrostate according to the optimization polarity as the positive class. For comparison with a pairwise connectivity baseline, we repeated the same procedure using the Fisher *z*-transformed within-*n*-plet functional connectivity (Section V-C) as the input feature instead of Ω. For each order *n*, dataset, and polarity, we plotted PR-AUC values against *n*-plet size, and reported these as order-dependent classification curves in Fig. 3e–f.

## Code availability

Code used to perform the analyses and generate the figures is available at https://github.com/camilo-espinosa/high-order-anaesthesia/.

## Data availability

For both multi-anaesthesia and DBS dataset, raw data are available for access from B.J. through academic collaboration.

## Supplementary Information

